# Dnmt3a1 regulates hippocampus-dependent memory via the downstream target Nrp1

**DOI:** 10.1101/2023.05.22.541739

**Authors:** Janina Kupke, Julien Klimmt, Franziska Mudlaff, Maximilian Schwab, Pavlo Lutsik, Christoph Plass, Carsten Sticht, Ana M.M. Oliveira

**Author notes:** Equal contribution (Julien Klimmt, Franziska Mudlaff). Ana M.M. Oliveira. Department of Molecular and Cellular Cognition Research, Central Institute of Mental Health, Medical Faculty Mannheim, Heidelberg University, 68159 Mannheim, Germany.

## Abstract

Epigenetic factors are well established players in memory formation. Specifically, DNA methylation is necessary for the formation of long-term memory in multiple brain regions including the hippocampus. Despite the demonstrated role for DNA methyltransferases (Dnmts) in memory formation, it is unclear whether individual Dnmts have unique or redundant functions in long-term memory formation. Furthermore, the downstream processes controlled by Dnmts during memory consolidation have not been investigated. In this study, we demonstrated that Dnmt3a1, the predominant Dnmt in the adult brain, is required for long-term spatial object recognition and contextual fear memory. Using RNA-sequencing, we identified an activity-regulated Dnmt3a1-dependent genomic program in which several genes were associated with functional and structural plasticity. Furthermore, we found that some of the identified genes are selectively dependent on Dnmt3a1, but not its isoform Dnmt3a2. Specifically, we identified Neuropilin 1 (Nrp1) as a downstream target of Dnmt3a1 and further demonstrated the involvement of Nrp1 in hippocampus-dependent memory formation. Importantly, we found that Dnmt3a1 regulates hippocampus-dependent memory via Nrp1. In contrast, Nrp1 overexpression did not rescue memory impairments triggered by reduced Dnmt3a2 levels. Taken together, our study uncovered a Dnmt3a-isoform-specific mechanism in memory formation, identified a novel regulator of memory, and further highlighted the complex and highly regulated functions of distinct epigenetic regulators in brain function.

## Introduction

Precise control of gene transcription is required for the functional and structural neuronal remodelling that underlies memory formation and maintenance(1–3). Gene transcription is orchestrated by the activity of transcription factors(4,5), epigenetic factors, and 3D-genomic factors such as chromatin structure or nuclear organization(6–9). Indeed, epigenetic mechanisms such as DNA methylation have emerged as important regulators of cognitive processes(8).

DNA methylation has traditionally been viewed as a static mechanism with important roles in transcription repression, cell fate determination, imprinting and silencing of transposons during early development (10). However, it is now well accepted that DNA methylation plays a critical role in the regulation of genomic responses in mature neurons. Several lines of evidence demonstrated dynamic and quick changes in DNA methylation in response to neuronal activity(11,12) and pharmacological and genetic studies established a causal link between DNA methyltransferase (Dnmt) activity and memory formation. Specifically, the *de novo* DNA methyltransferase, Dnmt3a, plays a documented function in cognitive abilities(13–17). Furthermore, it has recently been shown that *DNMT3A* variants are the underlying cause of the neurodevelopmental disorder Tatton-Brown-Rahman syndrome, which is characterized by intellectual disability(18,19). Despite this evidence, the downstream processes regulated by Dnmt3a that contribute to memory formation are unknown. Moreover, the genomic program regulated by Dnmt3a and required for the formation of long-term memory remains uncovered.

The Dnmt3a genomic locus gives rise to two isoforms, Dnmt3a1 and Dnmt3a2(20), that exhibit different genomic binding(21) and expression regulation in response to neuronal activity. Dnmt3a2 is transcribed from an intronic promoter and behaves like an immediate early gene, whereas the expression of Dnmt3a1 is not regulated at the level of transcription by neuronal activity(22–24). The requirement for Dnmt3a2 in memory formation has been demonstrated(23–25); however, the contribution of Dnmt3a1, despite its high abundance in the adult brain(26–28) has not been investigated. The differential regulation of expression of the two isoforms by neuronal activity and the studies that showed that normal Dnmt3a1 expression levels do not compensate the memory impairments found in Dnmt3a2 knockdown mice (23,24,29,30), suggest nonredundant functions by the two isoforms. However, whether the two enzymes are differentially required for and regulate distinct downstream processes during memory formation remains to be investigated.

In this study, we demonstrated that Dnmt3a1 is crucial for memory formation and for controlling an activity-regulated genomic program. We found that acute hippocampal reduction of Dnmt3a1 led to memory impairments in adult mice in the spatial object recognition test and contextual fear conditioning tasks without affecting hippocampus-independent tasks. Moreover, using RNA Sequencing analysis, we identified activity-regulated genes dependent on Dnmt3a1 levels. Intriguingly, we found that some of the identified genes are selectively dependent on Dnmt3a1, but not Dnmt3a2. Among these genes was Neuropilin-1 (Nrp1), a member of the semaphorin complex with an established function in synaptic plasticity(31,32). In a next step, we demonstrated the involvement of Nrp1 in long-term spatial object recognition and contextual fear memories. Furthermore, we showed that Dnmt3a1 regulates hippocampus-dependent memory via Nrp1. In contrast, Nrp1 overexpression did not rescue memory impairments triggered by reduced Dnmt3a2 levels, suggesting an isoform-specific mechanism for memory processes. Taken together these findings uncovered a Dnmt3a1-dependent genomic program and identified a downstream effector during memory formation.

## Materials and Methods

### Animals

3-month-old C57BL/6N male mice (Charles River, Sulzfeld, Germany) were used and housed 2-3 per cage (except during severe fighting) with *ad libitum* access to water and food, under a 12h light/dark cycle at 22 ± 1°C and 55 ± 10% relative humidity. Behavioural experiments were conducted during the light phase. Mice that were sick or injured were excluded. Uninjured cage mates were kept in the study. Performance exclusion criteria were applied to the spatial object recognition task as described below. Animals were randomly assigned to experimental groups and analysis was conducted blindly. Procedures complied with German animal care guidelines and European Community Council Directive 86/609/EEC.

### Recombinant Adeno Associated Virus (rAAV) production

Viral particles were produced and purified as described previously(33) with minor modifications. Specifically, AAV-293 cells (Stratagene #240073, California, USA) were used for co-transfection of target AAV plasmid and helper plasmids (pFΔ6, pRV1, and pH21). Viral particles in media were concentrated by incubation with 40% Polyethylene glycol (PEG) solution (pH 7,4) for 3h at 4°C and centrifugation. The viral pellet was added to lysed HEK cells prior to purification and concentration. For expression of shRNAs, we used a vector containing the U6 promoter upstream of the shRNA sequence and a chicken β-actin promoter to drive GFP expression. The sequences are as following: Dnmt3a1-shRNA1: GCAGACCAACATCGAATCCAT; Dnmt3a1-shRNA2: GGGAGGATGATCGAAAGGAAGGAGA; Dnmt3a2-shRNA: ACGGGCAGCTATTTACAGAGC; Nrp1-shRNA1: GGAAACCAAGAAGAAATATTA; Nrp1-shRNA2: GGGAGAGGAAATCGGAGCTAA; Control-shRNA: ACTACCGTTGTTATAGGTG. For temporally controlled knockdown of Dnmt3a1, we used a dual-component TetON-based system. The driver plasmid contained a neuron-specific promoter (hSynapsin) controlling the transactivator (rtTA), the tetracycline repressor (TetR) and Kusabira Orange (KO). The second construct contained GFP and a miR30-based shRNA targeting Dnmt3a1 controlled by the tetracycline-responsive promoter (TRE). This vector was generated using the plasmids pPRIME-CMV-GFP-FF3(34) (a gift from Stephen Elledge (Addgene plasmid # 11663; http://n2t.net/addgene:11663; RRID:Addgene_11663)) and pAAV-PTRE-tight-hM3Dg-mCherry(35) (a gift from William Wisden [Addgene plasmid # 66795; http://n2t.net/addgene:66795; RRID:Addgene_66795]). Specifically, the Dnmt3a1-shRNA1 sequence was inserted into the pPRIME vector and subsequently, the GFP-miR30-Dnmt3a1-shRNA expression cassette was subcloned into the vector containing the Tet-inducible promoter by replacement of the hM3Dq-mCherry insert. The overexpression of Nrp1-HA or a control gene, was achieved by placing HA-tagged Nrp1 or LacZ under the control of hSynapsin promoter, respectively. The infection rate, toxicity, viral titer, and knockdown efficiency for each batch of generated viruses were evaluated. The final titer was around 1-2 x10^12^ viral particles/ml.

### Fluorescent labelling

Sedated mice underwent intracardial perfusion with ice-cold PBS followed by 4% paraformaldehyde (PFA, Sigma-Aldrich, Munich, Germany). Brains were post-fixed overnight with 4% PFA solution at 4°C and subsequently transferred to 30% sucrose (in PBS). 30 μm slices were cut using a cryotome (CM1950, Leica), and incubated in Hoechst 33258 (2 μg/ml, Serva, Heidelberg, Germany) for 5 min. Slices were imaged using a 20x oil objective on a TCS SP8 confocal microscope (Leica Microsystems, Oberkochen, Germany).

### Stereotaxic surgery

rAAVs were injected into the dorsal hippocampus (dHPC) at the following coordinates relative to the bregma: −2 mm anteroposterior, ± 1.5 mm mediolateral, −1.7, −1.9, and −2.1 mm dorsoventral from bregma. Each spot received 500 nl of viral solution injected at a speed of 200 nl/min through a 33G needle. After injections at each site, the needle was left in place for 60 s. During behavioural experiments, the experimenter was blinded to the virus injected into each mouse. Behavioural testing commenced 3 weeks after rAAV delivery.

### Behavioural testing

Before behavioural testing, mice were habituated to the experimenter and behavioural room by handling for 3 consecutive days, 1 min/mouse. Different mouse cohorts were used for testing short-term (1h) or long-term memory (24h). The spatial object recognition (SOR) test and contextual fear conditioning (CFC) were performed as previously described(3). In SOR, mice were habituated (6 min) to an arena (50 cm × 50 cm × 50 cm) with a visual cue placed on the arena wall, followed by a training session that consisted of three 6 min-exposures (3 min intertrial interval) to two objects. In the testing session (1h or 24h later) one object was moved to a new location and object exploration was scored for 6 min. If in one experimental batch during the SOR test control animals did not show a preference, due to day effects, the whole set of animals was excluded from this analysis but was still included in contextual fear conditioning data. Further, an animal was excluded if total exploration time during testing was inferior to 2 seconds. One week later, CFC was performed in which mice were allowed to explore the conditioning chamber (23 × 23 × 35 cm, TSE, Bad Homburg, Germany) for 148 s followed by a 2 s foot shock (0.5 mA). They remained 30 s in the conditioning chamber before returning to their home cage (HC). During testing, mice were exposed 5 mins to the conditioning chamber. Where applicable, mice received intraperitoneal doxycycline hyclate (2.5 mg in 500 µL saline solution, 100mg/kg, Sigma-Aldrich, Munich, Germany) 3 days prior to CFC.

The accelerating rotarod and cued fear conditioning were performed in an independent animal cohort. The rotarod apparatus (Ugo Basile, Italy) had a 3-cm diameter rod with speed increasing from 4 to 40 r.p.m. over 5 min. Testing occurred 3 times daily with 1-hour intervals over 3 days. Trials began at the same time of the day and ended either with mice falling off or after 300 s. In cued fear conditioning, mice were put into a chamber (23 × 23 × 35 cm, TSE, Germany). A white noise cue (75dB) played from 2 to 2.5 minutes after placement, followed by a 0.7 mA foot shock in the last 2 seconds of the cue. Mice stayed in the chamber for an extra 30 seconds before returning to their cage. During testing, mice experienced a new context for 6 minutes (different chamber, smooth flat floor, altered dimensions, and a novel odor). The cue started 3 minutes after placing the mice in the chamber and lasted for 3 minutes.

### Primary hippocampal cultures and pharmacological treatments

Hippocampal cultures from newborn C57Bl/6N mice (Charles River, Sulzfeld, Germany) were prepared and maintained as previously described(3). In brief, mice hippocampi were dissociated at P0 by papain digestion and plated onto tissue culture dishes coated with poly-D-lysine and laminin (Sigma-Aldrich, Munich, Germany). The primary cultures were maintained for 8 days in Neurobasal-A medium (Gibco™) supplemented with 1% rat serum (Biowest), 0.5 mM L-glutamine (Sigma-Aldrich, Munich, Germany) and B27 (Gibco™), followed by incubation in transfection media: salt-glucose-glycine solution (10 mM HEPES, pH 7.4, 114 mM NaCl, 26.1 mM NaHCO3, 5.3 mM KCl, 1 mM MgCl2, 2 mM CaCl2, 30 mM glucose, 1 mM glycine, 0.5 mM C3H3NaO3, and 0.001% phenol red) and phosphate-free Eagle’s minimum essential medium (9:1 v/v), supplemented with insulin (7.5 μg/ml), transferrin (7.5 μg/ml), and sodium selenite (7.5 ng/ml) (ITS Liquid Media Supplement, Sigma-Aldrich, Munich, Germany) and penicillin-streptomycin. rAAV infection occurred on day in vitro (DIV) 4. Experiments were performed on DIV 9-11. To induce action potential bursting, cultures were treated with 50 µM bicuculline (Alexis Biochemicals, Farmingdale, NY, USA). DNA transfection was performed on DIV 8 using Invitrogen Lipofectamine 2000 (Invitrogen, CA, USA). The DNA (μg): Lipofectamine (μl) ratio was 1.6:5 in 1 ml of medium. The Lipofectamine/DNA mixture was left on the cells for 3 h before it was replaced with a transfection medium. Doxycycline hyclate (25 μM, Sigma-Aldrich, Munich, Germany) was introduced in the medium at DIV 8. To monitor doxycycline dependency, images of infected primary neurons with the TetOn-based miR30 system were acquired (Axio Vert.A1, Zeiss, Oberkochen, Germany).

### qRT-PCR primer design

The quantitative reverse-transcription PCR (qRT-PCR) primers were designed with Primer3 (https://primer3.ut.ee/) using either the RefSeq curated annotation or the GENCODE VM23 comprehensive transcript annotation, along with the GRCm38/mm10 mouse genome assembly. The specificity and amplicon product size of the primers were verified by BLAST search and *in silico* PCR (UCSC Genome Browser, mm10). Primer pair efficiencies and product melting curves were validated by qRT-PCR on serially diluted complementary DNA (cDNA) from primary mouse hippocampal cultures (refer to the “qRT-PCR” section). The list of all primers used in the study is provided in Sup Table 1.

### qRT-PCR

RNA from hippocampal cultures or tissue was extracted using the RNeasy Plus Mini Kit (Qiagen, Hilden, Germany) with extra DNase I digestion on the column, following the manufacturer’s instructions. For extraction from mouse tissue, the infected region (identified by GFP expression) was quickly dissected. RNA was transcribed into cDNA using the High-Capacity cDNA reverse-transcription kit (Applied Biosystems, Foster City, CA, USA). qRT-PCR was performed on Step One Plus Real-Time PCR System (Applied Biosystems, Foster City, CA, USA) using TaqMan gene expression assays (Applied Biosystems, Foster City, CA, USA) or the Power SYBR Green PCR Master Mix (Applied Biosystems) assays. The following TaqMan gene expression assays were used: Dnmt1 (Mm00599763_m1), Dnmt3a1 (Mm00432870_m1), Dnmt3a2 (Mm00463987_m1), Dnmt3b (Mm00599800_m1) and the housekeeping gene Gusb (Mm00446953_m1). Power SYBR Green PCR Master mix was used for the remaining target genes. PCR reactions were run as technical triplicates in 10 µL reactions (96-well format) using 0.5 μM of each primer. 2 μL of diluted cDNA (about 1.25 ng) was added to each reaction. The thermal cycling was conducted with the following settings: a 10-minute incubation at 95 °C, 40 cycles of 10 seconds each at 95 °C, 60 °C, and 72 °C, followed by a 15-second incubation at 95 °C. Melt curves were generated by heating from 60 °C to 90 °C at a ramp rate of 0.6 °C/min. Relative expression levels of each target transcript were determined by the ΔΔCt method using beta-Actin mRNA levels as a reference(36).

### Western Blotting

Primary hippocampal neurons infected on DIV 4 were lysed on DIV 10 in boiling SDS sample buffer (160 mM Tris-HCl (pH 6.8), 4% SDS, 30% glycerol, 10 mM dithiothreitol, and 0.02% bromophenol blue). In the case of western blotting of tissue samples, dHPC was quickly dissected in ice-cold PBS and homogenized in RIPA buffer (150 mM NaCl, 1% Triton X-100, 0.5% sodium deoxycholate, 0.1% SDS, 50 mM Tris, pH 8.0) supplemented with 1% protease inhibitor cocktail (Sigma-Aldrich, Munich, Germany). Protein concentration was measured by Bradford assay and 20 µg of denatured protein (in Laemmli buffer at 95°C for 5 min) was loaded in a 7.5% polyacrylamide gel and blotted onto a nitrocellulose membrane (GE Healthcare, Buckinghamshire, UK). The membranes were blocked in 5% milk in PBS with 0.01% Tween (PBST) and probed with the following antibodies (diluted in 5% milk in PBST) overnight at 4 °C: α-Tubulin (1:400000, Sigma t9026), α-Dnmt3a (1:5000, H-295, Santa Cruz, SC-20703), α-HA-tag (1:7500, Covance, MMS101R). The next day, membranes were incubated with horseradish peroxidase-conjugated secondary antibodies (1:5000 diluted in 5% milk in PBST) for 1h at room temperature. Finally, they were analysed using ChemiDoc^TM^ Imaging System (Bio-Rad, California, USA).

### Immunocytochemistry

Primary hippocampal neurons plated on coverslips were rinsed with PBS and fixed with a prewarmed solution of 4% PFA and 4% sucrose for 15 min at room temperature. After permeabilizing the cells in ice-cold methanol for 6 min, blocking was performed with 10% normal goat serum in PBS for 1 h at room temperature. Next, cells were incubated with primary antibody α-Dnmt3a (1:500, H-295, Santa Cruz, SC-20703diluted in PBS containing 2% BSA, 0,1% Triton X-100) overnight at 4 °C, which continued with secondary antibody incubation (1:500 goat anti-mouse Alexa488 [Life Technologies, Eugene, OR, USA]) diluted in PBS containing 2% BSA, 0,1% Triton X-100) for 1 h at room temperature. Finally, coverslips were treated with Hoechst 33258 (2 μg/ml, Serva, Heidelberg, Germany) for 5 min and mounted on glass slides. Images were acquired with a 40x oil objective on a fluorescence microscope (Leica Microsystems, Oberkochen, Germany).

### Bulk RNA Sequencing

Total RNA from mouse hippocampal cultures was isolated as above described and 500 ng of total RNA was used for bulk RNA-sequencing. Libraries were sequenced as 100bp paired-end reads, amounting to approximately 61 million reads per sample. Data processing was performed with R (version 3.6.3) and bioconductor (version 3.9) in Rstudio (version 1.1.463). Quality control of clean sequencing reads was performed using FastQC (Babraham Bioinformatics). Low-quality reads were removed using trim_galore (version 0.6.4). The resulting reads were aligned to the mouse genome version GRCm38.p6 and counted using kallisto version 0.46.1 (37). The count data were transformed to log2-counts per million (logCPM) using the voom-function from the limma package(38). Differential expression analysis was performed using the limma package in R. A false positive rate of α= 0.05 with FDR correction was taken as the level of significance. Volcano plots and heatmaps were created using ggplot2 package (version 2.2.1) and the complexHeatmap package (version 2.0.0)(39). For enrichment analysis, we used the fgsea(40), the enrichmentbrowser(41), and the Enrichr packages(42).

### Statistical analysis

Each data set was subjected to a normality test prior to further comparisons (Shapiro-Wilk normality test; alpha = 0.05). For normally distributed data sets, a two-tailed unpaired Student’s t-test was performed to compare two groups. If more than two groups were analyzed simultaneously, a one-way ANOVA was used followed by appropriate multiple comparison *post hoc* tests to control for multiple comparisons as specified. In case of a non-Gaussian distribution, two-tailed Mann-Whitney tests were used to compare two distinct groups, or a Kruskal-Wallis test followed by Dunńs *post hoc* test to compare more than two groups. The sample size was determined based on similar experiments carried out in the past and the literature. All plotted data represent mean ± SEM. Statistical analysis was performed using GraphPad Prism 7 (GraphPad Software). All behavioural sessions were video recorded and manually scored to determine the exploration of objects during training and testing phases or freezing behaviour by an experimenter blind to the group identity. The TSE Systems Fear Conditioning program was used to score the mean velocity during the training for CFC.

## Results

### Dnmt3a1 is required for long-term memory formation

In this study, we sought to investigate the role of DNA methyltransferase 3a1 (Dnmt3a1) in neuronal function and memory formation. We employed stereotaxic surgery and recombinant adeno-associated viruses (rAAVs) to deliver a control shRNA sequence (Control-shRNA)(3) or two independent shRNAs to knockdown Dnmt3a1 into the adult (8 weeks) dorsal hippocampus (dHPC) of mice (Figure 1A, B). We used the rAAV1/2 serotype given its well established neuronal tropism(43). To monitor viral expression upon infection, the viral constructs contained a GFP cassette under the control of the chicken-beta-actin promoter (Figure 1A, B). We confirmed knockdown efficiency of our shRNAs by transfection of primary hippocampal neurons *in vitro* and immunostaining against endogenous Dnmt3a1 (Sup Figure 1A). To further validate the knockdown efficiency *in vivo*, we performed qRT-PCR and western blot analysis of dHPC of mice infected with the viral constructs (Sup Figure 1B-G). We confirmed that both sequences selectively knockdown Dnmt3a1 without affecting other *Dnmts*. shRNA1 and shRNA2 sequences effectively reduced Dnmt3a1 expression at both the mRNA (Sup Figure 1B; two-tailed, unpaired t-test; *Dnmt3a1*: t(_6_)=6.28; *Dnmt3a2*: t(_6_)=0.48; *Dnmt1*: t(_6_)=0.81; *Dnmt3b*: t(_6_)=0.09; C; two-tailed, unpaired t-test; *Dnmt3a1*: t(_8_)=5.94; *Dnmt3a2*: t(_7_)=1.07; *Dnmt1*: t(_8_)=0.52; *Dnmt3b*: t(_8_)=1.17;) and protein (Sup Figure 1D-G; two-tailed, unpaired t-test; Dnmt3a1-shRNA1: t(_6_)=6.15; Dnmt3a1-shRNA2: t(_7_)=3.16) levels. To evaluate a requirement for Dnmt3a1 in memory performance, we conducted hippocampus-dependent memory tests on mice three weeks after stereotaxic surgery (Figure 1C). We found that, when tested 24 hours after training in the spatial object recognition test, mice injected with either Dnmt3a1-shRNA1 or-shRNA2 exhibited no preference for the displaced object (Figure 1D; two-tailed, unpaired t-test; Dnmt3a1-shRNA1: t(_17_)=2.43; Dnmt3a1-shRNA2: t(_22_)=3.15). However, when evaluated 1 hour after training, mice in shRNA groups explored the displaced object for a length of time comparable to the control group (Figure 1E; two-tailed, unpaired t-test; Dnmt3a1-shRNA1: t(_10_)=2.04; Dnmt3a1-shRNA2: t(_18_)=0.64). Importantly, Dnmt3a1 knockdown did not affect the overall amount of time that mice spent exploring the objects during training (Sup Figure 1H, I; two-tailed, unpaired t-test; Dnmt3a1-shRNA1: t(_17_)=0.92; Dnmt3a1-shRNA2: t(_22_)=0.35). To assess further the role of hippocampal Dnmt3a1 in memory formation, we trained mice in contextual fear conditioning. Knockdown of Dnmt3a1 with either shRNA1 or shRNA2 impaired contextual fear memory response 24 hours after training (Figure 1F; two-tailed, unpaired t-test; Dnmt3a1-shRNA1: t(_20_)=2.60; Dnmt3a1-shRNA2: t(_20_)=2.54) without decreasing freezing levels compared to the Control-shRNA when tested 1 hour after training (Figure 1G; two-tailed, unpaired t-test; Dnmt3a1-shRNA1: t(_15_)=0.04; Dnmt3a1-shRNA2: t(_22_)=0.90). This decrease in freezing was not due to altered responsiveness to the shock administration during the training session (Sup Figure 1 J, K; two-tailed, unpaired t-test; Dnmt3a1-shRNA1; Pre-Shock: t(_20_)=0.16; Shock: t(_20_)=0.25; Dnmt3a1-shRNA2: Pre-Shock: t(_21_)=1.06; Shock: t(_21_)=0.36). These results show that Dnmt3a1 knockdown impairs long-term memory formation without affecting short-term memory. Next, we investigated whether the memory deficits are specific to hippocampus-dependent tasks. We assessed the performance of mice injected with Control-shRNA, Dnmt3a1-shRNA-1 or Dnmt3a1-shRNA-2 in the auditory cue fear conditioning (Sup Figure 1L) and accelerating rotarod (Sup Figure 1M), two tasks that do not require hippocampal function(44,45). We found that hippocampal Dnmt3a1 knockdown did not affect cued long-term fear memory (Sup Figure 1L; one-way ANOVA test followed by Dunnett’s multiple comparisons test; F(_2,32_)=0.18, p=0.84) or motor skill learning (Sup Figure 1M; two-way repeated measures ANOVA; effect of training day F_(1.661,53.15)_=11.76; effect of virus F(_2,32_)=0.51), indicating region-specific deficits.

**Figure 1:**
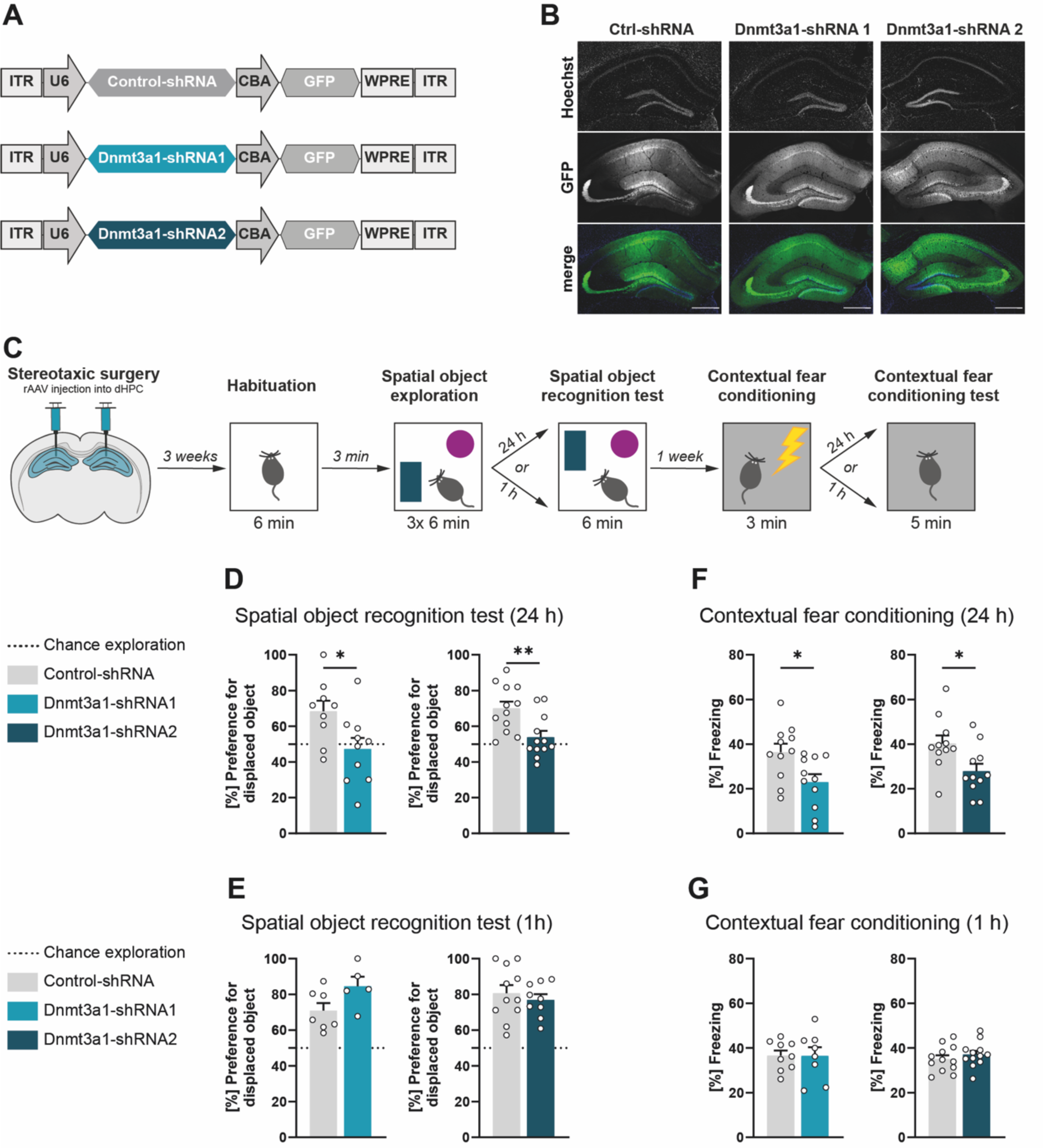
Reduced hippocampal Dnmt3a1 levels impair memory formation. (**A**) Schematic representation of viral constructs. (**B**) Representative images of the dorsal hippocampus of mice infected with Control-shRNA, Dnmt3a1-shRNA1 or Dnmt3a1-shRNA2. Scale bar: 100 µm. (**C**) Schematic representation of experimental design. (**D, E**) Long-term (D) or short-term (E) spatial object recognition memory of mice expressing Control-shRNA (n=7-12), Dnmt3a1-shRNA1 (n=5-10) or Dnmt3a1-shRNA2 (n=9-12). Dashed lines represent equal preference for either object (chance exploration). *p<0.05, **p<0.01 by two-tailed, unpaired t-test. (**F, G**) Long-term (F) or short-term (G) contextual fear memory of mice expressing Control-shRNA (n=9-11), Dnmt3a1-shRNA1 (n=8-11) or Dnmt3a1-shRNA2 (n=11-13). *p<0.05 by two-tailed, unpaired t-test.

The behavioural testing in this series of experiments was carried out three weeks following stereotaxic delivery of rAAVs to achieve high viral expression and, as a result, high knockdown efficiency. However, the persistent reduction of Dnmt3a1 levels may lead to network-wide changes that could impact learning and memory. Consequently, in order to more directly assign a role for Dnmt3a1 in mature neurons during the process of memory formation, we adopted a temporally controlled method that limited Dnmt3a1 knockdown to the time of behavioural testing. We used a TetON-based system that consisted of two components: for tight control of Tet-dependent transgene expression in neurons, one viral construct contained the human synapsin (hSyn) promoter controlling the expression of the reverse tetracycline-controlled transactivator (rtTA) together with the Tet repressor (TetR) and the fluorescent protein Kusabira Orange (KO) that serves as an infection marker(46,47). In the second construct, the Dnmt3a1-specific shRNA sequence 1 or the Control-shRNA was embedded in a microRNA-30-based expression cassette(34), which is under the control of the tetracycline responsive promoter (TRE) (Figure 2A-B). In the absence of doxycycline, TetR is bound to TRE inhibiting its activity. Upon doxycycline administration, TetR releases from the TRE promoter allowing the binding of the activator rtTA. Administration of doxycycline to hippocampal primary cultures verified tight control of transgene expression via the Tet-On system (Sup Figure 2 A-D). The expression of GFP after doxycycline administration increased over a time course of 72 hours (Sup Figure 2B). Western blot analysis of Dnmt3a1 at the same time points revealed a significant knockdown of Dnmt3a1 protein levels at 72 hours (Sup Figure 2C, D; two-tailed, unpaired t-test; 0h: t(_10_)=1.03; 48h: t(_10_)=1.86; 72h: t(_10_)=3.06; Mann-Whitney test; 24h: U=10). Thus, doxycycline was injected intraperitoneally 72 hours before behavioural testing (Figure 2C). We found that knocking down hippocampal Dnmt3a1 around the time of fear conditioning reduced freezing levels when mice were tested 24 hours after training (Figure 2D; two-tailed, unpaired t-test: t(_27_)=2.99); however, mice froze to a similar extent than the control mice when testing occurred 1 hour after training (Figure 2E; two-tailed, unpaired t-test: t(_18_)=0.32). The responsiveness to the shock administration during the training session did not cause this decrease in freezing (Sup Figure 2E; two-tailed, unpaired t-test; Pre-Shock: t(_14_)=0.47; Shock: t(_14_)=0.80). Taken together, these findings further support a function of Dnmt3a1 during long-term memory formation and suggest that Dnmt3a1 participates in the regulation of learning-dependent gene expression required for long-term memory.

**Figure 2:**
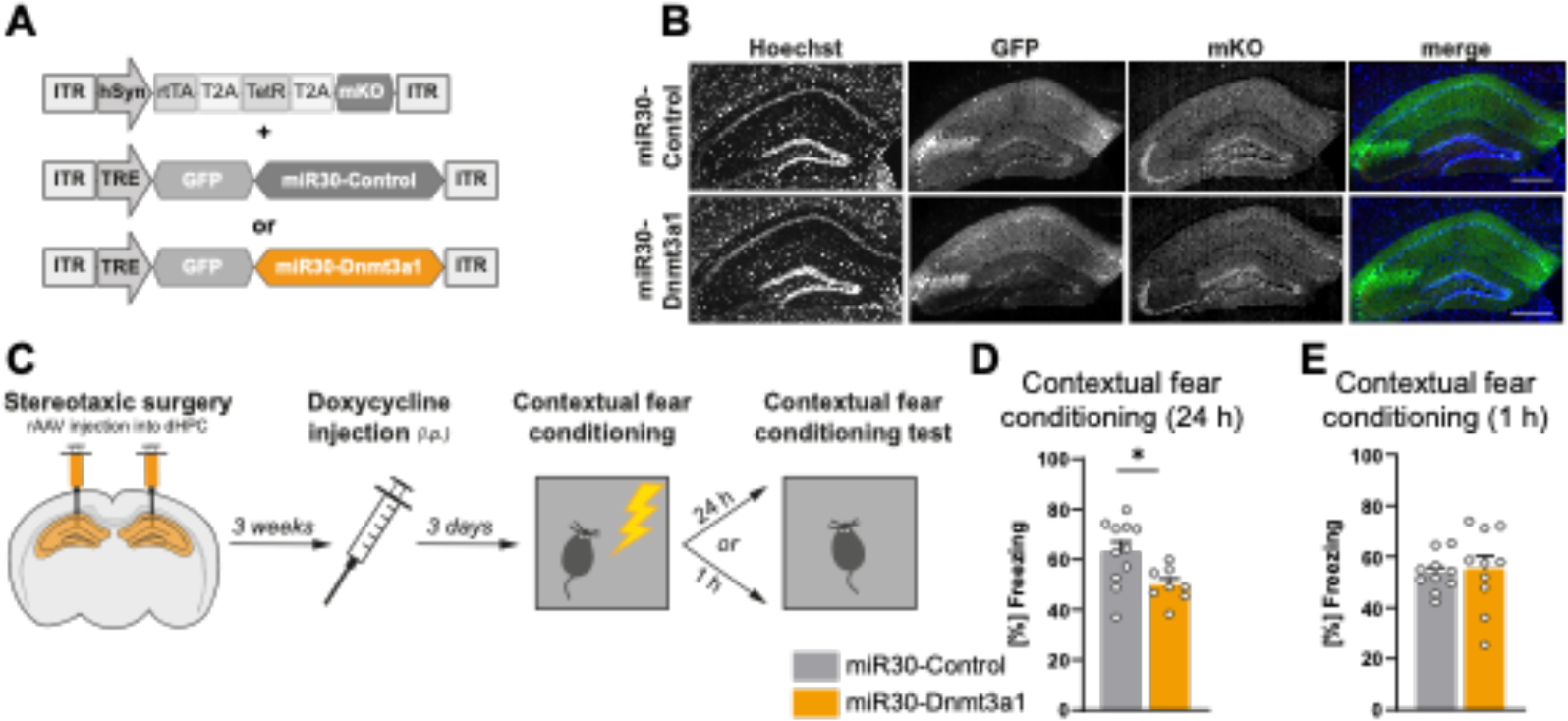
Temporally-restricted Dnmt3a1 knockdown impairs memory consolidation. (**A**)Schematic representation of the viral constructs encoding the TetON-based system. The driver construct expresses under the hSyn promoter the Tet repressor (TetR), the reverse tetracycline-controlled transactivator (rtTA) and the infection marker Kusabira Orange (mKO). The Tet response element (TRE)-dependent construct contains the TRE promoter driving the miR30-based shRNA cassettes. In the absence of doxycycline, TRE is inhibited by TetR which results in suppression of the miR30 system. In the presence of doxycycline, TetR loses its affinity thus enabling rtTA to bind to TRE and activate the expression of the miR30 system. (**B**) Representative images of the dHPC of mice infected with rAAVs expressing the miR30-Control shRNA or miR30-Dnmt3a1-shRNA2 sequence. Scale bar: 100 µm. (**C**) Schematic representation of experimental design. (**D, E**) Long-term (D) or short-term (E) contextual fear memory of mice expressing miR30-Control shRNA (n=12-17) or miR30-Dnmt3a1 shRNA (n=10-14). *p<0.05, by two-tailed, unpaired t-test.

### Dnmt3a1 regulates the expression of activity-dependent genes involved in synaptic plasticity

Next, to elucidate the molecular mechanism by which Dnmt3a1 regulates memory consolidation we sought to identify activity-dependent genes regulated by Dnmt3a1. To this end, we performed RNA sequencing of hippocampal neurons infected with Dnmt3a1-shRNA2 or a Control-shRNA after inducing neuronal activity by bicuculline treatment (Figure 3A, B, Sup Figure 3A, B). In control conditions, we found that neuronal activity leads to the downregulation of 1981 genes and upregulation of 1935 genes including established activity-regulated genes, e.g., Arc, Fos, Npas4 (Figure 3C, Sup File 1).

**Figure 3:**
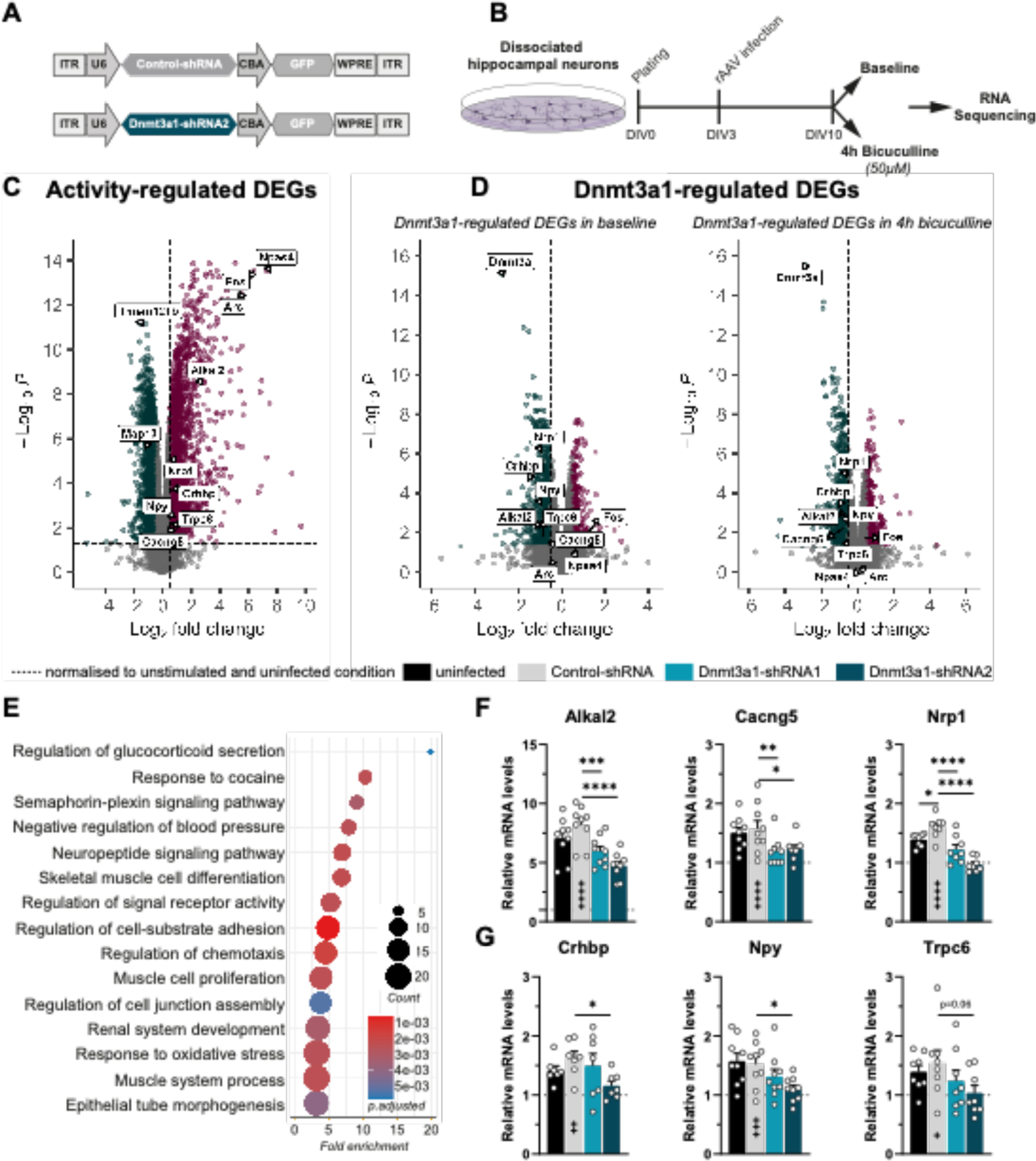
Dnmt3a1 regulates transcription of genes involved in synaptic plasticity processes. (**A**) Schematic representation of viral constructs. (**B**) Experimental design used to identify activity-regulated genes whose expression is altered upon Dnmt3a1 reduction. DIV: Day *in-vitro*. (**C**) Volcano plot of differentially expressed genes (DEGs) in control condition (infected with rAAVs expressing Control-shRNA) in response to neuronal activity via Bicuculline treatment; n=4 independent neuronal preparation per condition. Log2 fold change cut-off: ± 0.5; adjusted p-value cut-off: 0.05. (**D**) Volcano plot of DEGs upon infection with Control-shRNA or Dnmt3a1-shRNA2 in baseline condition or in stimulated condition (4h bicuculline); n=4 independent neuronal preparation per condition. Log2 fold change cut-off: ± 0.5; adjusted p-value cut-off: 0.05. (**E**) GO-Term analysis of overlapping genes between activity-regulated and Dnmt3a1-regulated DEGs. Dot plot illustrates Top 15 GO term enrichment of biological processes. (**F**) qRT-PCR analysis of *Alkal2*, *Cacng5*, *Nrp1* expression levels in hippocampal cultures infected with Control–shRNA or Dnmt3a1-shRNA1 or-shRNA2 and stimulated 4h with Bicuculline. Expression levels were normalised to the uninfected control in baseline conditions (dashed line); (n=8-9 independent neuronal cultures). *p<0.05, ***p≤0.001, ****p≤0.0001 by one-way ANOVA test followed by Sidak’s multiple comparisons test(**G**) qRT-PCR analysis of *Crhbp*, *Npy*, *Trpc6* expression levels in hippocampal-cultured cells infected with Control–shRNA or Dnmt3a1-shRNA1 or-shRNA2 and stimulated 4h with Bicuculline. Expression levels were normalised to the uninfected control in baseline conditions (dashed line); (n=7-9 independent neuronal cultures). *p<0.05, **p≤0.01, ns: not significant by one-way ANOVA test followed by Sidak’s multiple comparisons test. Control-shRNA 4h bicuculline versus unstimulated: ^+^p<0.05, ^++^p<0.01, ^+++^p≤0.001, ^++++^p≤0.0001 by one-way ANOVA test followed by Sidak’s multiple comparisons test.

To identify genes dependent on Dnmt3a1, we focused on genes that are differentially expressed in Dnmt3a1 knockdown versus control conditions in basal (Figure 3D, Sup File 2) and/or upon bicuculline treatment (Figure 3D, Sup File 2). Notably, the expression of classical immediate early genes, such as Arc, Fos, Npas4 was not altered in Dnmt3a1-shRNA conditions (Figure 3D, Sup File 2), suggesting that the identified changes are not a result of an overall disruption of neuronal responses. We hypothesised that Dnmt3a1-regulated genes that are involved in memory formation should be a) differentially expressed in response to Dnmt3a1 reduction and b) activity-regulated. Thus, we overlapped the activity-regulated (Figure 3C, Sup File 1) and the Dnmt3a1-regulated gene sets (Figure 3D, Sup File 2) in order to find potential activity-regulated Dnmt3a1 target genes. This intersection identified 491 candidates (Sup Figure 3B,C, Sup File 3). Gene ontology (GO) analysis of the 491 genes yielded a strong enrichment for terms related to structural and functional plasticity, namely *“Neuropeptide signaling pathway”*, *“Regulation of signaling receptor activity”* and *“Regulation of cell-substrate adhesion”* (Figure 3E, Sup File 3). This suggests that genes downstream of Dnmt3a1 may play an important role in synaptic plasticity and learning and memory. We confirmed some of the identified Dnmt3a1 target genes in independent biological samples using qRT-PCR (Figure 3F, G). To further rule out the possibility of off-target effects, this validation was performed using the two independent shRNA sequences. We confirmed that the expression of *Alkal2*, *Cacng5*, *Nrp1, Crhbp, Npy and Trpc6* is activity-regulated (Figure 3F,G; one-way ANOVA test followed by Sidak’s multiple comparisons test: *Alkal2*: F(_6,56_)=78.46, p<0.0001; *Cacng5*: F(_6,56_)=16.67, p<0.001; *Nrp1*: F(_6,49_)=46.53, p<0.0001; *Crhbp*: F(_6,42_)=13.60, p<0.0001; *Npy*: F(_6,56_)=15.66, p<0.001; *Trpc6*: F(_6,49_)=5.373, p=0.003). The expression of the Control-shRNA sequence appeared to cause a mild effect on the expression of Nrp1, but not of the other genes (comparison to uninfected control). However, this was not replicated across independent experiments (see Sup Figure 3F). Furthermore, we found that the expression of *Alkal2*, *Cacng5*, and *Nrp1* is reduced upon Dnmt3a1 depletion achieved by shRNA1 or shRNA2 (Figure 3F), whereas *Crhbp, Npy and Trpc6* failed to show a significant effect with one of the shRNA sequences (Figure 3G).

*Dnmt3a* encodes for two distinct isoforms Dnmt3a1 and Dnmt3a2, which are differentially regulated by neuronal activity(22,23). We further investigated whether the two isoforms regulate distinct targets. To test this, we infected primary hippocampal cultures with a previously validated shRNA to reduce Dnmt3a2 levels(23,24) (Sup Figure 3D) and investigated the impact of this manipulation on the expression of the identified Dnmt3a1-regulated genes. Reduction of Dnmt3a2 did not change the expression of *Alkal2*, *Cacng5* or *Nrp1* neither in basal conditions nor upon neuronal activity (Sup Figure 3E; one-way ANOVA test followed by Sidak’s multiple comparisons test: *Alkal2*: F(_5,25_)=49.04, p<0.0001; *Cacng5*: F_(_(_5,25_)=4.96, p=0.0044; *Nrp1*: F(_4,25_)= 4.96, p=0.0044). Taken together these findings demonstrate that Dnmt3a1 is required for the regulation of a synaptic plasticity-related transcriptional program and further suggest that the Dnmt3a2 isoform may regulate a distinct gene pool.

### Neuropilin-1 is a Dnmt3a1 downstream target required for memory formation

Next, we aimed at investigating whether Dnmt3a1 role in memory consolidation requires the activity of identified target genes. Among the genes whose expression is controlled by Dnmt3a1, we focused on neuronal genes which have been identified to be regulated by novel environment exposure in the mouse hippocampus(48). These included, among others, the previously validated gene Neuropilin-1 (*Nrp1)* (Figure 3G, H). Nrp1 is known to build a complex with Semaphorin 3a (Sema3A) and Plexin A4 and is implicated in signalling pathways regulating neuronal morphology(31,49). Although it has been demonstrated that this complex is important for AMPA receptor trafficking(50), is still unknown whether Nrp1 plays a role in hippocampus-dependent memory formation. We performed a shRNA-mediated loss-of-function approach to assess the role of Nrp1 in memory formation. We generated two independent shRNA constructs to reduce the expression of *Nrp1* (Figure 4A) and validated their knockdown efficiency *in vivo* after stereotaxic delivery of the viral constructs or a Control-shRNA into the dHPC. We confirmed that Nrp1-shRNA1 and -shRNA2 sequences led to a robust reduction of Nrp1 mRNA levels (Sup Figure 4A, B; two-tailed, unpaired t-test: Nrp1-shRNA1: t(_10_)=5.28; Nrp1-shRNA2: t(_16_)=7.25). Mice were subjected to a hippocampus-dependent behavioural memory task three weeks after viral delivery (Figure 4B). Knockdown of Nrp1 with either shRNA resulted in exploration of the displaced object at chance levels (Figure 4C; one-way ANOVA test followed by Dunnett’s multiple comparisons test: F(_2,29_)=4.46, p=0.0204), as well as in decreased freezing responses in the contextual fear conditioning test compared to mice injected with the Control-shRNA (Figure 4D; one-way ANOVA test followed by Dunnett’s multiple comparisons test: F(_2,30_)=6.13, p=0.0059). Notably, Nrp1 knockdown did not affect the overall object exploration time (Sup Figure 4C; one-way ANOVA test followed by Dunnett’s multiple comparisons test: F(_2,32_)=0.80, p=0.4584) or the responsiveness to the shock administration (Sup Figure 4D; one-way ANOVA test followed by Sidak’s multiple comparisons test: F(_5,62_)=83.97, p<0.0001). In conclusion, this set of experiments uncovered a novel function for Nrp1 in long-term memory formation.

**Figure 4:**
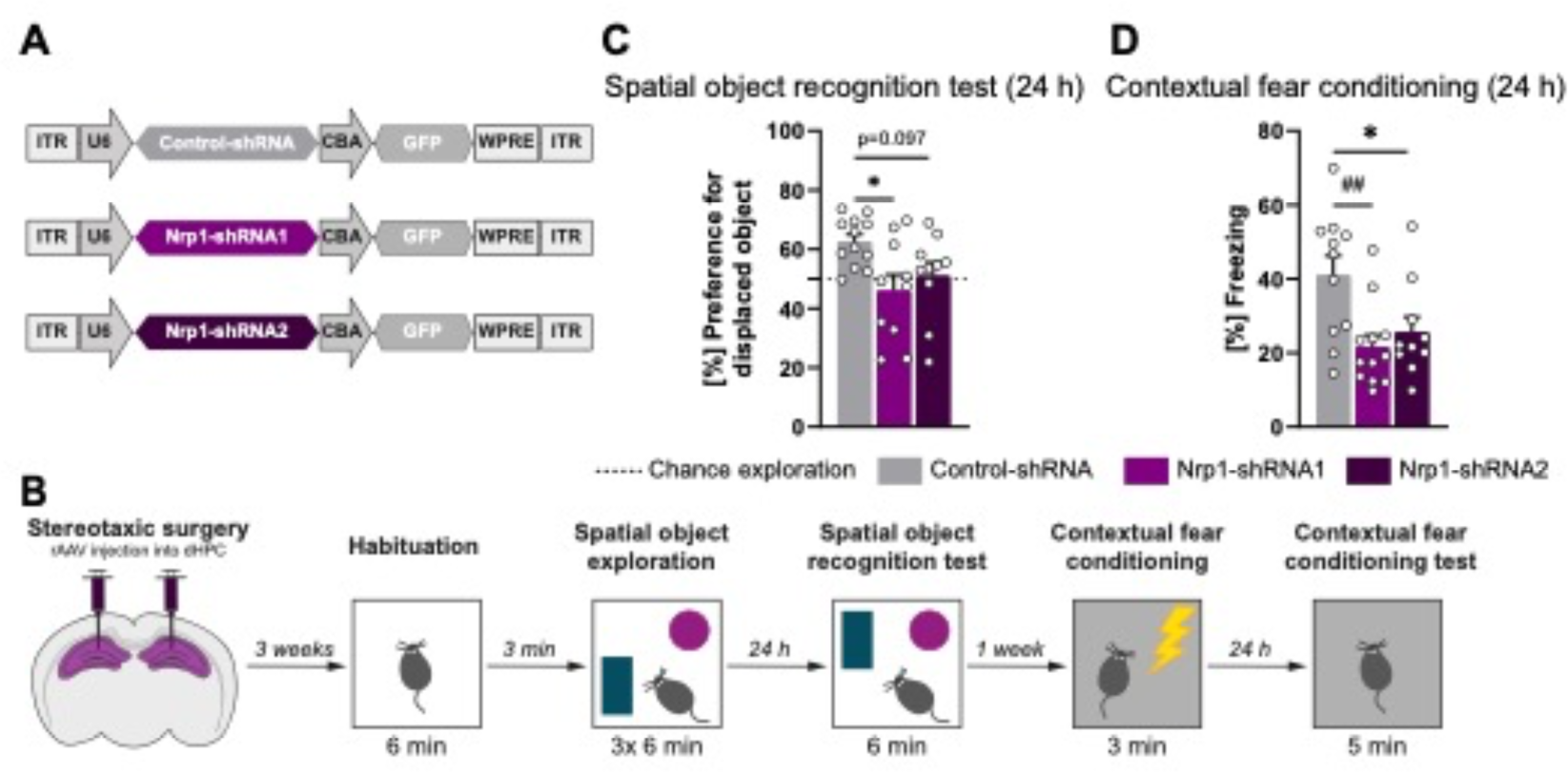
Hippocampal Neuropilin-1 is required for memory formation. (**A**) Schematic representation of viral constructs. (**B**) Schematic representation of experimental design. (**C**) Long-term spatial object recognition memory of mice expressing Control-shRNA (n=12), Nrp1-shRNA1 (n=10) or Nrp1-shRNA2 (n=10). Dashed lines represent equal preference for either object (chance exploration). *p<0.05 by one-way ANOVA test followed by Dunnett’s multiple comparisons test. (**D**) Long-term contextual fear memory of mice infected with rAAVs expressing Control-shRNA (n=11), Nrp1-shRNA1 (n=12) or Nrp11-shRNA2 (n=10). *p<0.05 by one-way ANOVA test followed by Dunnett’s multiple comparisons test. ^##^p≤0.01 by Kruskal-Wallis test followed by Dunn’s multiple comparisons test.

### Dnmt3a1, but not Dnmt3a2, regulates long-term memory formation via its downstream target Nrp1

To evaluate a function of Nrp1 as a downstream effector of Dnmt3a1 during memory formation, we performed a rescue experiment. To this end, we overexpressed HA-tagged Nrp1 or a control protein (LacZ) under the regulation of hSyn promoter together with either Dnmt3a1-shRNA2 or Control-shRNA sequences in the mouse hippocampus (Figure 5A, B). We first confirmed successful overexpression of Nrp1 in the dHPC by western blot analysis (Sup Figure 5A, B). After stereotaxic delivery of the constructs, mice underwent the spatial object recognition task (Figure 5B). The different groups did not exhibit differences in the time spent exploring the objects during the training session (Sup Figure 5C; one-way ANOVA test followed by Dunnett’s multiple comparisons test: F(_3,48_)=0.12, p=0.9477). As shown above, the reduction of Dnmt3a1 levels caused memory impairments which are manifested by the decreased preference for the displaced object (Figure 1D). Strikingly, this impairment was rescued upon Nrp1 overexpression (Figure 5C; one-way ANOVA test followed by Sidak’s multiple comparisons test: F(_3,46_)=7.39, p=0.004). Mice that overexpressed Nrp1 and expressed a control shRNA sequence showed memory performance similar to the control group (Figure 5C). Taken together, these results demonstrate that Nrp1 acts downstream of Dnmt3a1, is required for memory formation and can per se rescue Dnmt3a1-dependent memory impairments. Thus, indicating that Dnmt3a1 regulates memory formation, at least in part, via its downstream target Nrp1.

**Figure 5:**
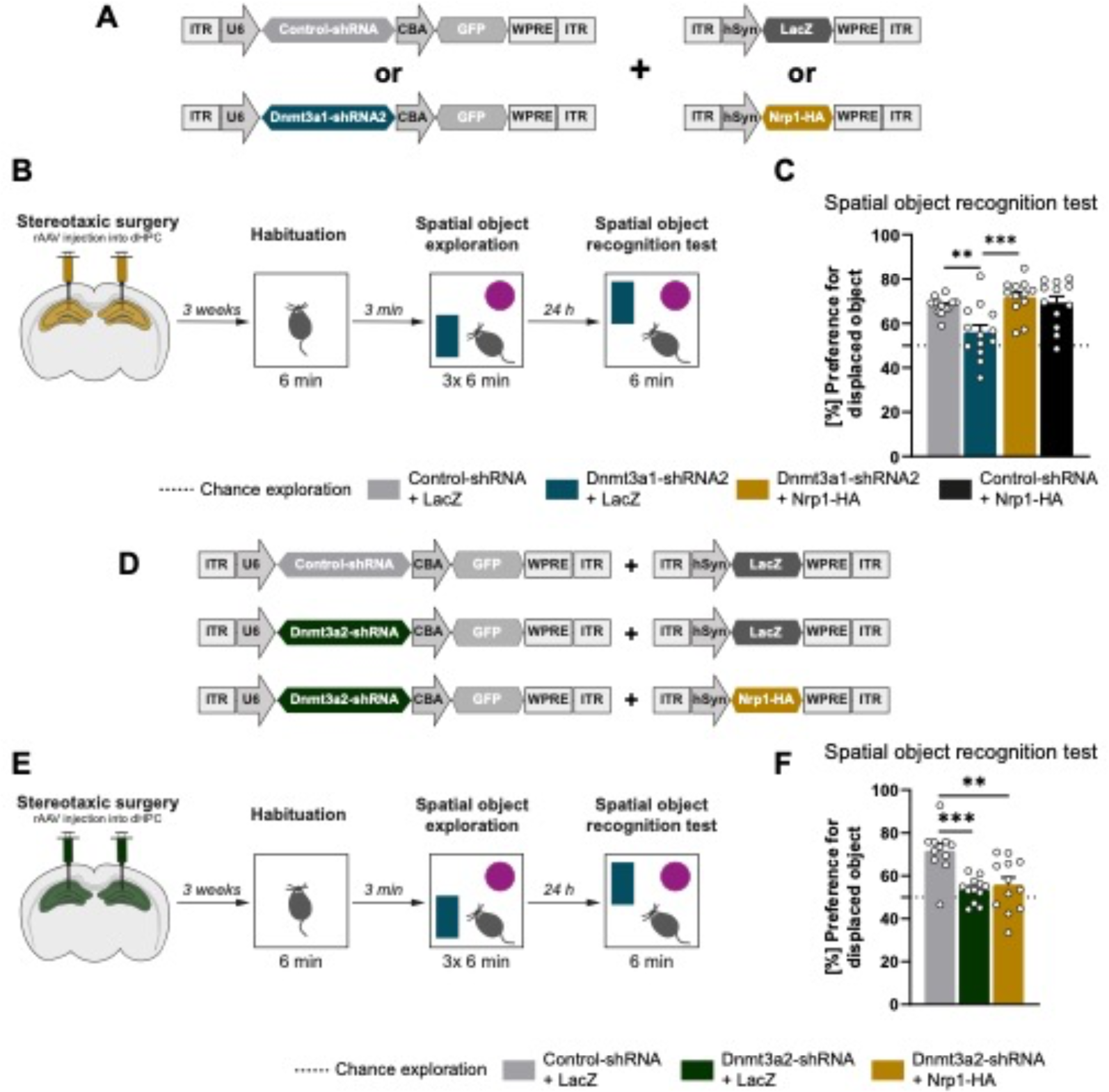
Neuropilin-1 rescues Dnmt3a1-, but not Dnmt3a2-, dependent memory impairments. (**A**) Schematic representation of viral constructs. (**B**) Schematic representation of experimental design. (**C**) Long-term spatial object recognition memory of mice expressing Control-shRNA + LacZ (n=13), Dnmt3a1-shRNA2 + LacZ (n=13), Dnmt3a1-shRNA2 + Nrp1-HA (n=12) or Control-shRNA + Nrp1-HA (n=12). Dashed lines represent equal preference for either object (chance exploration). **p≤0.01, ***p≤0.001 by one-way ANOVA test followed by Sidak’s multiple comparisons test. (**D**) Schematic representation of viral constructs. (**E**) Schematic representation of experimental design. (**F**) Long-term spatial object recognition memory mice expressing Control-shRNA + LacZ (n=11), Dnmt3a2-shRNA + LacZ (n=12) or Dnmt3a2-shRNA + Nrp1-HA (n=12). Dashed lines represent equal preference for either object (chance exploration). **p≤0.01, ***p≤0.001 by one-way ANOVA test followed by Sidak’s multiple comparisons test.

Our gene expression analysis suggested that the isoforms Dnmt3a1 and Dnmt3a2 regulate distinct genes. Specifically, we showed that Nrp1 expression is regulated by Dnmt3a1, but it is not altered upon knockdown of Dnmt3a2 (Sup Figure 3D). Next, to investigate a specific functional link between Dnmt3a1, Nrp1 and memory formation, we tested if Nrp1 overexpression rescues Dnmt3a2-knockdown-dependent memory impairments. We overexpressed HA-tagged Nrp1 with a Dnmt3a2-specific shRNA sequence in the mouse dorsal hippocampus and performed the spatial object recognition task in mice (Figure 5D, E). Importantly, none of the conditions showed differences in overall time exploring the objects during training (Sup Figure 5D; one-way ANOVA test followed by Sidak’s multiple comparisons test: F(_2,32_)=1.92, p=0.1633). As previously described(23), knockdown of Dnmt3a2 impaired spatial object recognition memory and remarkably, this impairment was not rescued upon Nrp1 overexpression (Figure 5F; one-way ANOVA test followed by Sidak’s multiple comparisons test: F(_2,32_)=10, p=0.0002). This finding shows that the rescue obtained in response to Nrp1 overexpression is not a result of a generalized memory enhancement effect and further supports that Nrp1 acts downstream of Dnmt3a1 but not Dnmt3a2. Thus, this data uncovers a *Dnmt3a* isoform specific mechanism in memory formation.

## Discussion

In this study we demonstrated a critical role for Dnmt3a1 in long-term memory formation by showing that acute knockdown of Dnmt3a1 in the adult hippocampus selectively impairs long-term memory formation and further identified activity-dependent genes regulated by Dnmt3a1 levels. Furthermore, our data suggests that despite a requirement for both Dnmt3a1 and Dnmt3a2 for memory formation, the two isoforms regulate this process via distinct mechanisms; Nrp1 overexpression rescued Dnmt3a1, but not Dnmt3a2, knockdown-driven impairments in memory formation. Taken together these findings have advanced our understanding of the requirement for distinct Dnmts in mnemonic processes as well as of the identity of downstream effector molecules.

Most studies that aimed at investigating the role of Dnmt3a in adult brain cognition have utilized heterozygous constitutive(51) or conditional knock-out mice(15,16,52). These models delete *Dnmt3a* during critical prenatal and postnatal neurodevelopmental phases, which may confound the interpretation of the function of *Dnmt3a*-coded proteins in memory formation in the adult. Mitchnick and colleagues (2015) used small interfering RNAs to acutely disrupt *Dnmt3a* expression in the adult hippocampus and provided compelling evidence for a role of *Dnmt3a*-coding genes in long-term hippocampal memory formation(14). In addition, our present and previous studies(23–25) have corroborated these findings and demonstrated that both Dnmt3a isoforms, Dnmt3a1 and Dnmt3a2, are required for memory formation in adult mature neurons. This study has further identified for the first time an activity-regulated genomic program regulated by Dnmt3a1 levels in hippocampal neurons. We found that the knockdown of Dnmt3a1 altered the expression of several genes both in baseline as well as upon neuronal activation. Even if counterintuitive at first, the reduction of Dnmt expression resulted not only in de-repression of gene expression, but also downregulation. Although these gene expression changes are likely a combination of direct effects on the altered genes and indirect mechanisms via the disruption of expression of other transcriptional regulatory processes, these findings corroborate previous studies in which a link between transcription activation and DNA methylation has been shown(10,28,53). In our transcriptomic analysis we found that Dnmt3a1 regulates the expression of several genes with key roles in synaptic plasticity and memory. Namely, we found changes in the expression of *Npy*, *Cort*, *Trpc6*, which have been implicated in memory consolidation(54–56). Upon the confirmation with two independent shRNA sequences, we focused on Nrp1 which has been shown to be regulated by enriched environment exposure(57) and within dentate gyrus memory engrams(48). Besides the established function for the semaphorins and their receptors (Nrp and plexin family members) in neurodevelopment(32,58), only a few lines of evidence demonstrated their involvement in homeostatic and Hebbian forms of plasticity in the adult hippocampus(31,50,59,60). Notably, Nrp1 is localised at synapses of the adult rat hippocampus(60) and secreted semaphorins acting on the sema-Nrp-plexin complex modulate synaptic connectivity in granule cells of dentate gyrus and pyramidal neurons of CA1(60) as well as AMPA receptor trafficking in CA3-CA1 synapses(50). The present study adds to this body of literature by demonstrating that Nrp1 is required for the formation of hippocampal long-term memory. Intriguingly, we showed that overexpression of hippocampal Nrp1 rescued Dnmt3a1 knockdown-dependent memory impairments. Thus, our findings suggest that the regulation of Nrp1-dependent mechanisms during memory consolidation is a mechanism via which Dnmt3a1 contributes to memory formation.

This study uncovered a dissociation in the downstream processes regulated by the Dnmt3a isoforms during hippocampal long-term memory formation. Notably, we showed that Nrp1 overexpression selectively rescued memory impairments promoted by the reduction of Dnmt3a1, but not Dnmt3a2, levels in the adult hippocampus. Our previous work had already demonstrated differential regulation of the two enzymes upon neuronal activity and learning. In contrast to Dnmt3a1, Dnmt3a2 mRNA levels are induced in response to action potential bursting and learning(23), cocaine administration(30) and an inflammatory response(29) in different regions of the central nervous system. Despite the lack of regulation at the level of transcription, Dnmt3a1 protein levels appear to be regulated upon neuronal activity(22). It has also been shown that Dnmt3a isoforms target unique genomic loci(21,61). Collectively, these findings indicate distinct regulatory functions. Specifically, during the formation of memory, lack of redundancy is further supported by our studies that showed that impairments in long-term memory formation promoted by the acute, reduced hippocampal expression of Dnmt3a1 or Dnmt3a2 are not compensated by the other isoform. Thus, the emerging picture is that the distinct Dnmts are required for cognitive abilities through the regulation of complementary but non-overlapping processes. The recruitment of Dnmts to DNA is thought to be guided by chromatin marks, namely histone modifications(62) and non-coding RNA species that direct gene-specific methylation patterns(63–67). Notably, differential DNA recruitment between Dnmt3a1 and Dnmt3a2 may be at least in part driven by distinct recognition of histone tail marks(21,28,68,69). Moreover, the N-terminus unique to Dnmt3a1 has been shown to confer specific chromatin recognition properties to this isoform(28,69). Future studies should elucidate the mechanisms that dictate the regulation of distinct downstream targets for Dnmt3a1 and Dnmt3a2 in response to neuronal activity.

Our findings highlight the complex and highly regulated role of distinct epigenetic regulators in brain function. The relevance of investigating DNA methylation processes in the nervous system is further underscored by the multiple lines of evidence that demonstrated that DNA methylation dysregulation underlies several pathological conditions, such as neurodevelopmental(70) and neurodegenerative(71) diseases as well as psychiatric conditions(72,73) and chronic pain(74). Therefore, the elucidation of the regulatory mechanisms controlled by the various epigenetic factors is required to gain further understanding of the pathophysiological mechanisms underlying these conditions.

## Acknowledgements

We thank David V.C. Brito, Kübra Gülmez Karaca and Stefanos Loizou for their critical comments on the manuscript. Celia García-Vilela, Carmen Leibold, Stefanos Loizou, and Benjamin Zeuch for technical support. hSyn-LacZ construct was kindly gifted by Prof. Dr. Daniela Mauceri. This work was supported by the Deutsche Forschungsgemeinschaft (DFG) [grant numbers OL 437/1, OL 437/2, OL 437/3 and OL 437/4 to A.M.M.O.], the Chica and Heinz Schaller Foundation [fellowship and research award to A.M.M.O.], and the Joachim Herz Stiftung [Add-on Fellowships for Interdisciplinary Life Science to J.Kupke.].

## Author Contributions

JKupke, AMMO designed research; JKupke, JKlimmt, FM, MS, AMMO performed experiments; JKupke, JKlimmt, FM, MS, CS, AMMO, PL, CP analyzed and interpreted data; JKupke, AMMO wrote the paper. All authors edited the manuscript.

## Data availability

The RNA-Sequencing data analysed in this study is publicly available at the National Center for Biotechnology Information (NCBI) Gene Expression Omnibus (GEO) with the accession number GSE232630.

