## Supplemental Figures for "Dnmt3a1 regulates hippocampus-dependent memory via the downstream target Nrp1"

[2]Present address: Department of Molecular and Cellular Neurobiology, Center for Neurogenomics and Cognitive Research, Vrije Universiteit Amsterdam, 1081 HV, Netherlands

[3]Present address: Institute for Stroke and Dementia Research, University Hospital, Ludwig-Maximilians-University Munich, 81377 Munich, Germany

[4]Present address: Integrated Program in Neuroscience, McGill University, Montreal, QC H3A 2B4, Canada; Centre for Research in Neuroscience, Research Institute of the McGill University Health Centre, Montreal QC, H3G 1A4, Canada

[5]Present address: Department of Medical Oncology, National Center for Tumor Diseases (NCT) Heidelberg, University Hospital Heidelberg, 69120 Heidelberg, Germany

[#]equal contribution

**Sup. Table 1. List of primers used for qRT-PCR.**

| Name | | Sequence |
|---|---|---|
| **Actin** | 5' Primer | TATCCTGACCCTGAAGTACC |
| | 3' Primer | CTCGGTGAGCAGCACAGGG |
| **Alkal2** | 5' Primer | TGAAACATCTCACAGGTCCTCT |
| | 3' Primer | GTGCAGTCTCTCGTGTTGTG |
| **Cacng5** | 5' Primer | CCCTCGTCAGCCTCTTCTTC |
| | 3' Primer | ACCACAAGAGAGAGGCCAGA |
| **Crhbp** | 5' Primer | CTGAAGGTATTTGATGGTTGGA |
| | 3' Primer | TCTTCATAGTGGGCAGAGGG |
| **Nrp1** | 5' Primer | GCAAGACTCGAATCCTCCC |
| | 3' Primer | CCAATGTGAGGGCCAACTTC |
| **Npy** | 5' Primer | CGTGTGTTTGGGCATTCTGG |
| | 3' Primer | TGCCATATCTCTGTCTGGTGA |
| **Trpc6** | 5' Primer | CCAGCTTCCGGGGTAATGAA |
| | 3' Primer | ACATGTATGCTGGTCCTCGA |

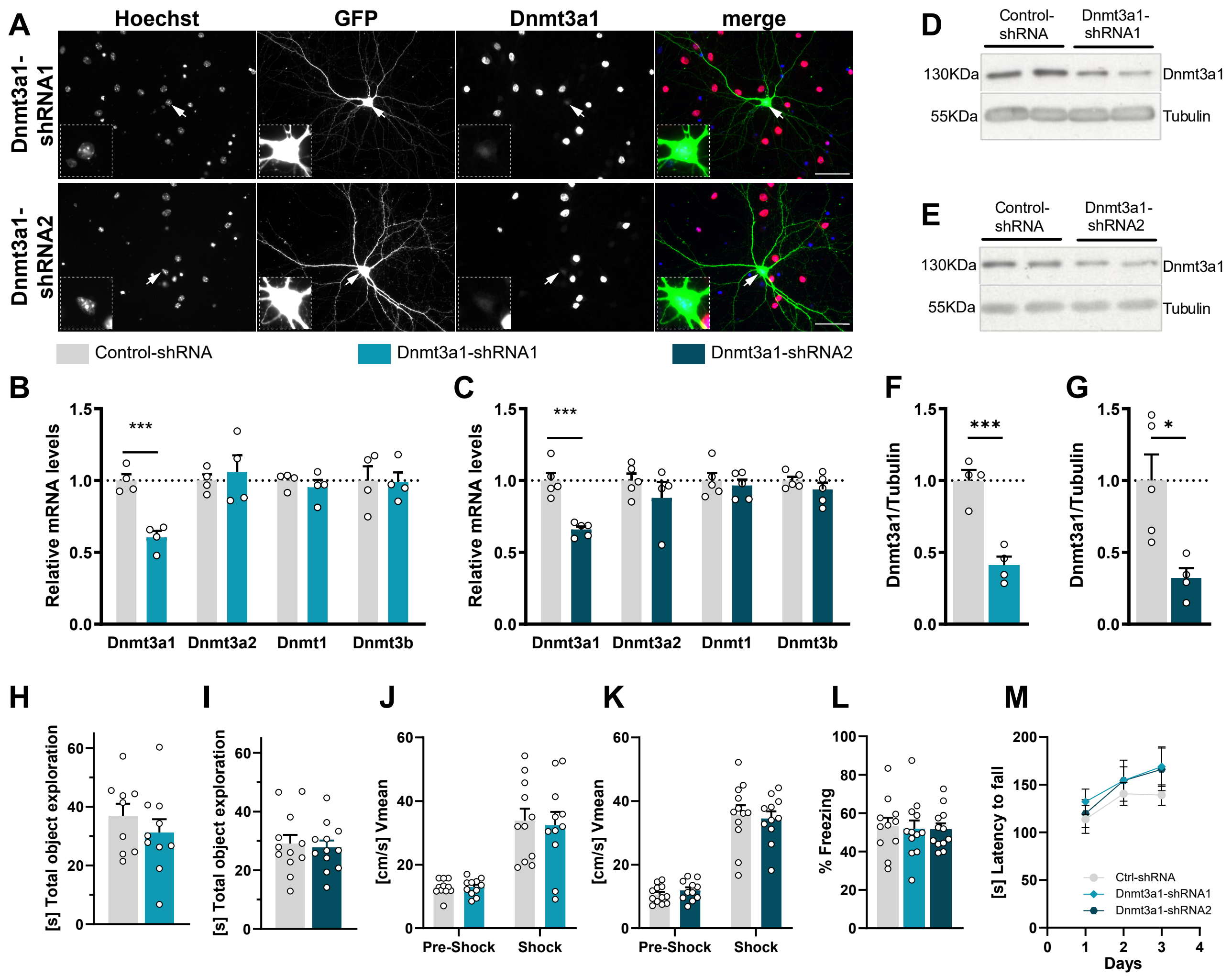


**Sup. Figure 1: Validation of Dnmt3a1 shRNA-dependent knock down and behavioural control parameters.** (**A**) Representative image of primary hippocampal cultures transfected with Dnmt3a1-shRNA1 or Dnmt3a1-shRNA2 (green) and immunostained against endogenous Dnmt3a levels (red). Scale bar: 50 µm. (**B, C**) Quantitative reverse-transcription PCR (qRT-PCR) analysis of the expression of DNA methyltransferases (Dnmt) in the dHPC of mice infected with rAAVs against Control-shRNA (n=4-5) or either (B) Dnmt3a1-shRNA1 (n=4) or (C) Dnmt3a1-shRNA2 (n=4). Dashed lines represent normalised expression levels to Control-shRNA. ***p≤0.001 by two-tailed, unpaired t-test. (**D, E**) Representative immunoblot scans of the expression of Dnmt3a1 or Tubulin in the dHPC of mice infected with rAAVs against Control-shRNA or either (D) Dnmt3a1-shRNA1 or (E) Dnmt3a1-shRNA2. (**F, G**) Immunoblot quantification of Dnmt3a1 protein levels in the dHPC of mice infected with rAAVs against Control-shRNA (n= 4-5) or either (F) Dnmt3a1-shRNA1 (n=4) or (G) Dnmt3a1-shRNA2 (n=4). Dashed lines represent normalised Dnmt3a1 to Tubulin expression of control group. *p<0.05, ***p≤0.001 by two-tailed, unpaired t-test. (**H, I**) Total object exploration time of mice infected with rAAVs against Control-shRNA (n=9-12) or either (H) Dnmt3a1-shRNA1 (n=10) or (I) Dnmt3a1-shRNA2 (n=12) during the three sessions of spatial object recognition training. No significant changes by two-tailed, unpaired t-test. (**J, K**) Mean velocities during CFC training pre-shock (Pre-Shock) or during shock (Shock) administration of the mice expressing rAAVs against Control-shRNA (n=11-13) or either (J) Dnmt3a1-shRNA1 (n=11) or (K) Dnmt3a1-shRNA2 (n=11). No significant changes by two-tailed, unpaired t-test. (**L**) Long-term cued fear memory of mice expressing Control-shRNA (n=11), Dnmt3a1-shRNA1 (n=12) or Dnmt3a1-shRNA2 (n=12). No significant changes by one-way ANOVA test followed by Dunnett's multiple comparisons test. (**M**) Motor skill learning of mice expressing Control-shRNA (n=11), Dnmt3a1-shRNA1 (n=12) or Dnmt3a1-shRNA2 (n=12). Effect of training day: p<0.001; effect of virus: p=ns by two-way repeated measures ANOVA. Ns= no significant.

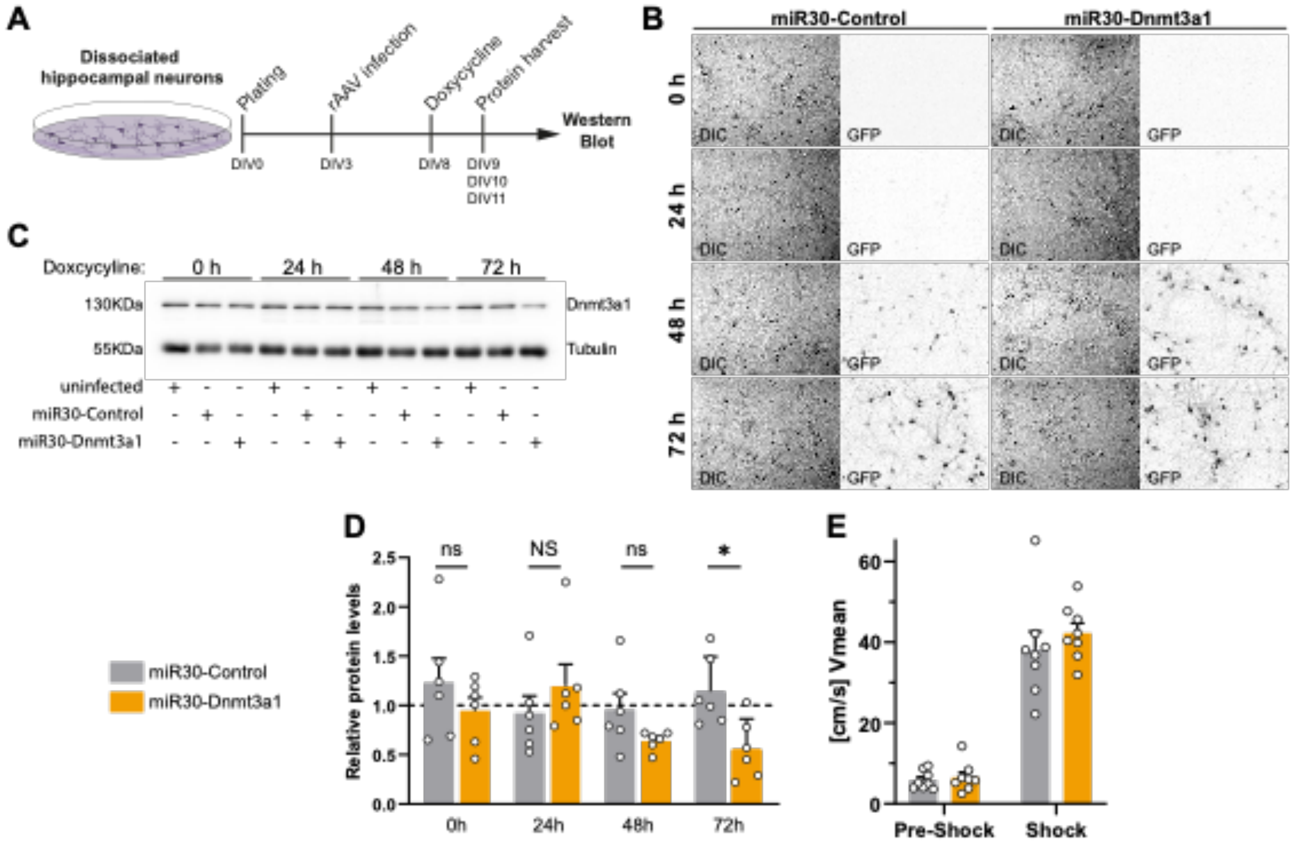


**Sup. Figure 2: Validation of mir30-based Dnmt3a1 shRNA-dependent knock down and behavioural control parameters.** (**A**) Experimental design used to identify the time-point of knock-down using the TetON-based miR30 system. DIV: Day *in-vitro*. (**B**) Representative images of primary hippocampal cells infected with miR30-Control or miR30-Dnmt3a1-shRNA2 sequence. Cultures were treated with doxycycline for 0h, 24h, 48, or 72h. (**C**) Representative immunoblot scans of the expression of Dnmt3a1 and Tubulin of primary hippocampal cultures infected with miR30-Control or miR30- Dnmt3a1-shRNA2 sequence and treated with doxycycline for either 0h, 24h, 48h or 72h. (**D**) Immunoblot quantification of Dnmt3a1 protein levels of primary hippocampal neurons infected with miR30-Control or miR30- Dnmt3a1-shRNA2 sequence and treated with doxycycline for either 0h, 24h, 48h or 72h. Expression levels were normalised to the uninfected controls (dashed line); (n=6 independent neuronal cultures). *$p \leq 0.05$, ns: not significant by two-tailed, unpaired t-test. NS: not significant by Mann-Whitney test. (**E**) Mean velocities during CFC training pre-shock (Pre-Shock) or during shock (Shock) administration of the mice expressing miR30-Control (n=8) or miR30-Dnmt3a1 (n=8) shRNAs. No significant changes by two-tailed, unpaired t-test.

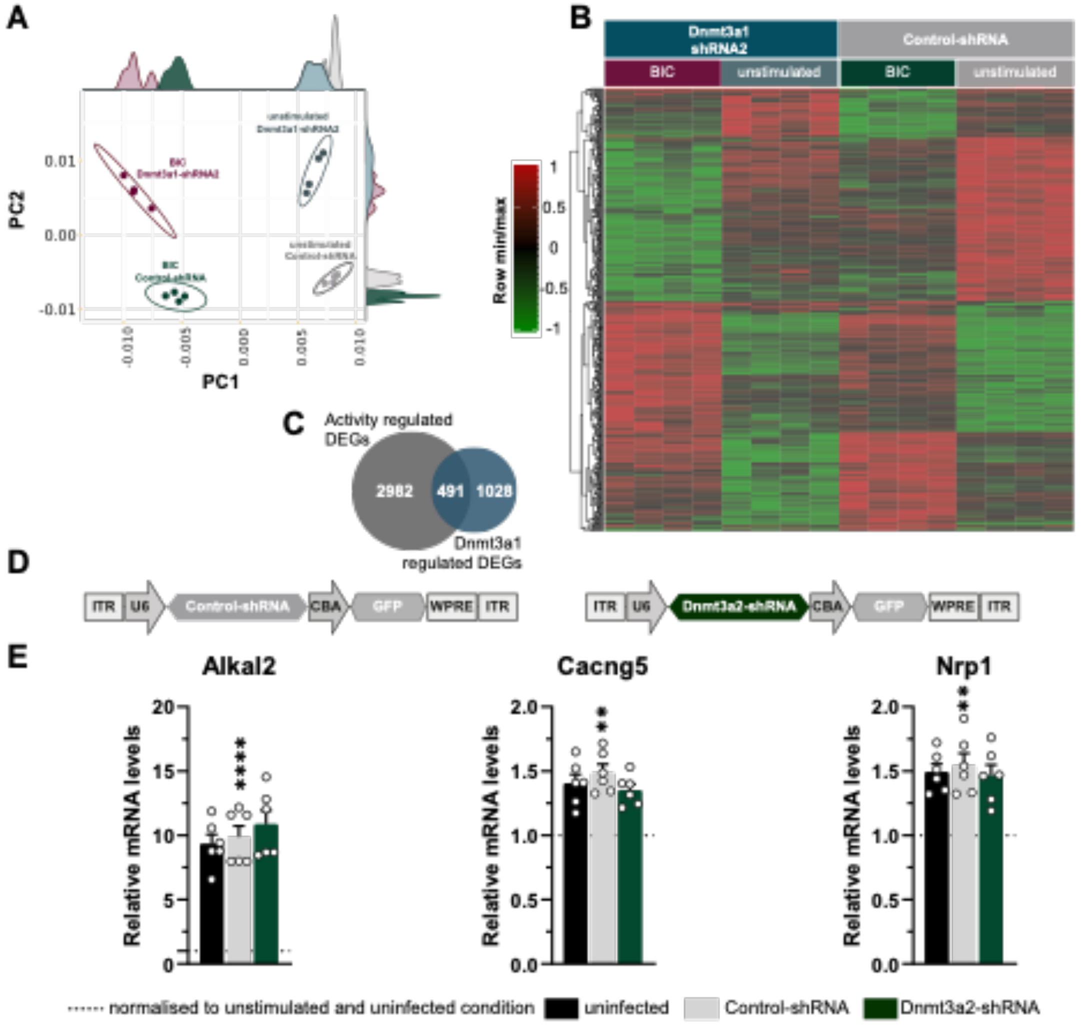


**Sup. Figure 3: PCA analysis and heat map of RNA sequencing data and effect of Dnm3a2 levels on the expression of selected genes.** (**A**) Principle-component analysis (PCA) of DEGs in primary hippocampal cultures infected with Control-shRNA or Dnmt3a1-shRNA2, that were either stimulated with Bicuculline for 4h or unstimulated; (n=4 independent neuronal culture preparations). (**B**) Heatmap of 491 activity-regulated Dnmt3a1-dependent genes (overlap between Figure 3C with Figure 3D) (n=4 independent neuronal cell preparations). Hippocampal cultures infected with rAAVs expressing Control-shRNA or Dnmt3a1-shRNA2, that were either stimulated with Bicuculline for 4h or unstimulated. (**C**) Venn Diagramm of activity regulated DEGs (Figure 3C) and Dnmt3a1 regulated DEGs (Figure 3D). (**D**) Schematic representation of viral constructs. (**E**) qRT-PCR analysis of Alkal2, Cacng5 and Nrp1 expression levels in hippocampal-cultured cells infected with Control–shRNA or Dnmt3a2-shRNA and stimulated 4h with Bicuculline. Expression levels were normalised to the uninfected controls in baseline conditions (dashed line); (n=6 independent neuronal cultures). Control-shRNA 4h bicuculline versus unstimulated: $**p<0.01$, $****p\leq0.0001$ by one-way ANOVA test followed by Sidak's multiple comparisons test. No significant differences between Control-shRNA and Dnmt3a2-shRNA by two-tailed, unpaired t-test.

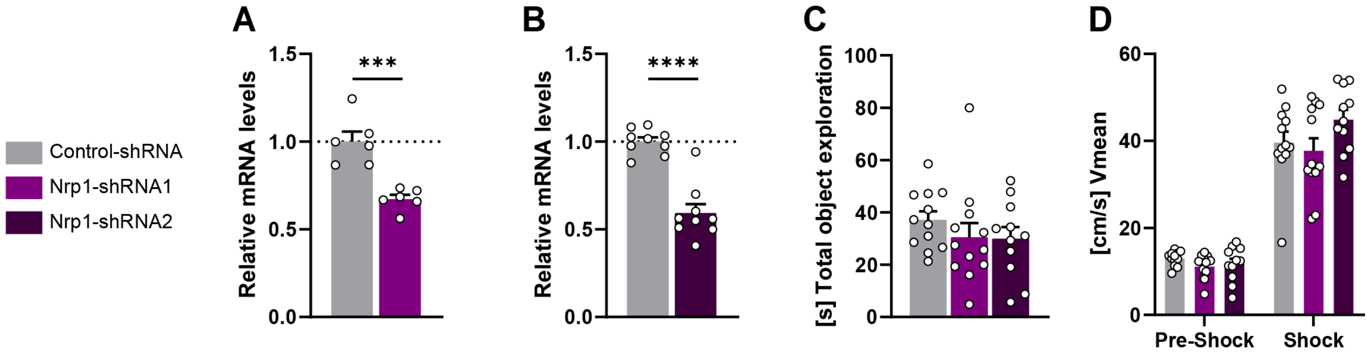


**Sup. Figure 4: Validation of Nrp1 shRNA-dependent knock down and behavioural control parameters.** (**A, B**) qRT-PCR analysis of Nrp1 expression in dHPC of mice infected with rAAVs against Control-shRNA (n=6-9), (A) Nrp1-shRNA1 (n=6) or (B) Nrp11-shRNA2 (n=9). Dashed lines represent normalised expression levels to Control-shRNA. ***p≤0.001, ***p≤0.0001, by two-tailed, unpaired t-test. (**C**) Total object exploration time of mice infected with rAAVs against either Control-shRNA (n=12) or Nrp1-shRNA1 (n=12) or Nrp1-shRNA2 (n=11) during the three sessions of spatial object recognition training. No significant changes between Control shRNA and either Nrp1-shRNA1 or Nrp1-shRNA2 by one-way ANOVA test followed by Dunnett's multiple comparisons test. (**D**) Mean velocities during CFC training pre-shock (Pre-Shock) or during shock (Shock) administration of the mice infected with either Control-shRNA (n=12) or Nrp1-shRNA1 (n=12) or Nrp1-shRNA2 (n=11). No significant changes between Control shRNA and either Nrp1-shRNA1 or Nrp1-shRNA2 by one-way ANOVA test followed by Sidaks multiple comparisons test.

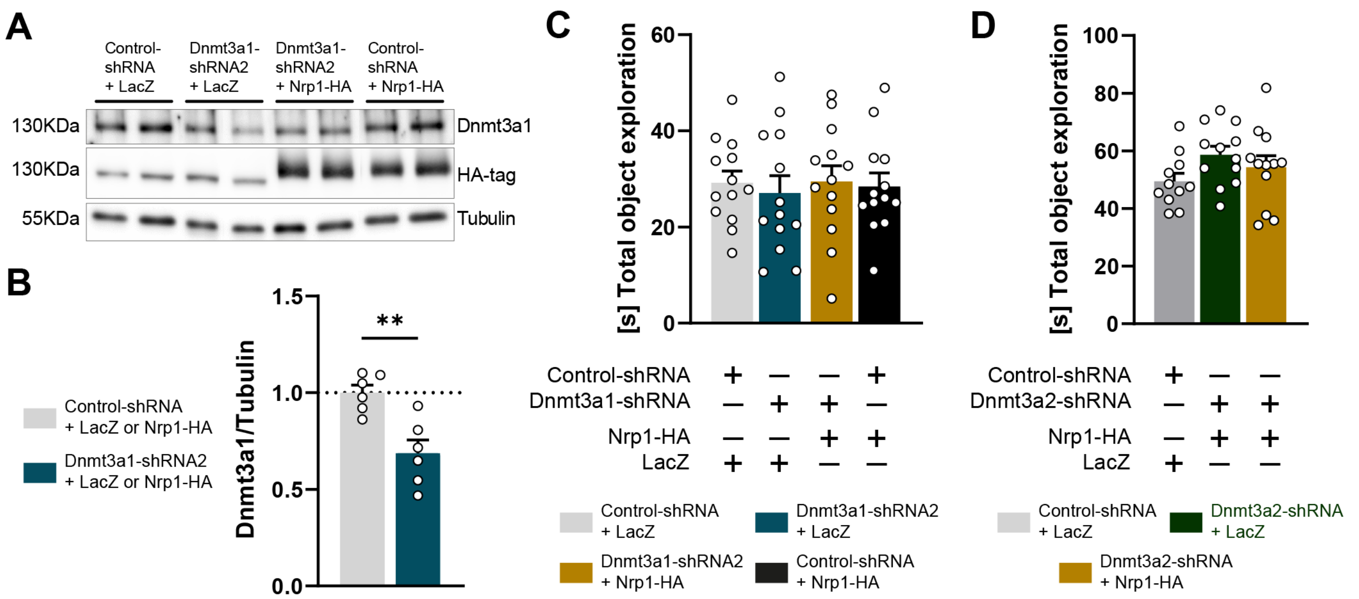


**Sup. Figure 5: Validation of Nrp1 overexpression and behavioural control parameters.** (**A**) Representative immunoblot scans of the expression of Dnmt3a1, HA-tagged Nrp1 or HA-tagged LacZ and Tubulin in the dHPC of mice infected with rAAVs as indicated. (**B**) Immunoblot quantification of Dnmt3a1 protein levels in the dHPC of mice infected with rAAVs against Control-shRNA (n= 6) or Dnmt3a1-shRNA2 (n=6). Dashed lines represent normalised Dnmt3a1 to Tubulin expression of control group. **p≤0.01 by two-tailed, unpaired t-test. (**C**) Total object exploration time of mice infected with Control-shRNA + LacZ (n=13), Dnmt3a1-shRNA2 + LacZ (n=13), Dnmt3a1-shRNA2 + Nrp1-HA (n=13) or Control-shRNA + Nrp1-HA (n=13) during the three sessions of spatial object recognition training. No significant changes by one-way ANOVA test followed by Sidak's multiple comparisons test. (**D**) Total object exploration time of mice infected with Control-shRNA + LacZ (n=11), Dnmt3a2-shRNA + LacZ (n=12), Dnmt3a2-shRNA + Nrp1-HA (n=12) during three sessions of spatial object recognition training. No significant changes by one-way ANOVA test followed by Dunnett's multiple comparisons test.
